# Emergent properties in microbiome networks reveal the anthropogenic disturbance of farming practices in vineyard soil fungal communities

**DOI:** 10.1101/2020.03.12.983650

**Authors:** Rüdiger Ortiz-Álvarez, Hector Ortega-Arranz, Vicente J. Ontiveros, Charles Ravarani, Alberto Acedo, Ignacio Belda

## Abstract

Agro-ecosystems are human-managed natural systems, and therefore are subject to generalized ecological rules. A deeper understanding of the factors impacting on the biotic component of ecosystem stability is needed for promoting the sustainability and productivity of global agriculture. Here we propose a method to determine ecological emergent properties through the inference of network properties in local microbial communities, and to use them as biomarkers of the anthropogenic impact of different farming practices on vineyard soil ecosystem functioning. In a dataset of 350 vineyard soil samples from USA and Spain we observed that fungal communities ranged from random to small-world network arrangements with differential levels of niche specialization. Some of the network properties studied were strongly correlated, defining patterns of ecological emergent properties that are influenced by the intensification level of the crop management. Low-intervention practices (from organic to biodynamic approaches) promoted densely clustered networks, describing an equilibrium state based on mixed (generalist-collaborative) communities. Contrary, in conventionally managed vineyards, we observed highly modular (niche-specialized) low clustered communities, supported by a higher degree of selection (more co-exclusion proportion). We also found that, although geographic factors can explain the different fungal community arrangements in both countries, the relationship between network properties in local fungal communities better capture the impact of farming practices regardless of the location. Thus, we hypothesize that local network properties can be globally used to evaluate the effect of ecosystem disturbances in crops, but also in when evaluating the effect of clinical interventions or to compare microbiomes of healthy vs. disturbed conditions.

## Introduction

Biota on Earth interacts. Organisms from the entire tree of life have different types of relationships with one another to ensure survival through genetic evolution and adaptation. These interactions have consequences for the whole ecosystem where these organisms thrive. Ecological communities can be defined by properties that result from the prediction of constituent taxa, properties known as Community Aggregated Traits (CATs)^1,2^. However, ecosystems can also be defined not through their constituent taxa, but from the emergent properties (EPs) that arise from specific community arrangements^3,4^. Emergent properties are directly related to the functionality of plant communities (i.e, seed survival rate), animal communities (i.e., animal behaviour; human societal interactions) and microbial communities (i.e, biofilm density, as a cause of composition behaviour), or all together as a whole (i.e., competition, predation strength). Both aggregated traits and emergent properties are characteristics that directly determine ecological processes, which further determine species pools or trophic fluxes ^5^.

Currently, most studies, particularly in microbial ecology, focus on correlative evidence between specific taxa abundance or diversity metrics and environmental factors or community phenotypes, as an attempt to understand the underlying mechanisms and resulting ecological processes^6^. Although valuable, it has been argued that this strategy is incomplete to understand the underlying mechanisms by which communities actually perform a function or process^5^. We argue that through the contextualization of emergent properties into ecological mechanisms, it would be possible to make predictions of how communities would behave under concrete circumstances. Additionally, translating those idiosyncratic community behaviours into a measurable metric can be a critical step for future microbiome monitoring applications, such as in sustainable farming^7^, food production or human health.

Smart farming harbours a demand of new biomarkers of soil health (see USDA definition at: https://www.nrcs.usda.gov/wps/portal/nrcs/main/soils/health/), since comprehensive information is often inaccessible to land managers^8^ and a single universal methodology to measure soil quality based on the microbiome does not exist yet^9,10^, despite notable efforts^11,12^. In particular, developing a strategy to mechanistically understand the fungal component of the soil microbiome, has implications in monitoring management of crops in risk of drought (i.e. vineyards or olive trees), since fungal-based food webs adapt better to drought than bacterial-based food-webs^13,14^. In this context, inferring emergent properties that relate to microbial functional processes and community stability becomes a priority of global interest. Unfortunately, these emergent properties are typically hard to measure, particularly in microbial ecology, given the high variety of microbes that are still being unveiled and contextualized into potential functional roles ^15,16^.

Here we aimed to implement new biomarkers of ecological disturbance based on ecological emergent properties by combining metacommunity theory and co-occurrence networks, regardless of the knowledge available on the different taxa comprising the metacommunity. A metacommunity is defined as a group of communities within the same habitat/region/pool that usually display multiple possible arrangements according to environmental filters, dispersal restrictions, priority effects and the latter established interactions ^17^. Co-occurrence networks derived from a metacommunity, which comprises the whole number of potential associations between all the taxa in the pool^18^, have been adequately used to understand taxa affinities to distinct ecological niches or geographic clusters ^19^. However, these hardly give an idea of the variety of potential arrangements of local communities, since each local community likely comprises only a fraction of taxa. We argue that merging the metacommunity-inferred associations into each of the local communities, will allow the estimation of network properties in all the local communities within the metacommunity, individually, obtaining local information on microbial ecosystem functioning **(Box 1)**. Yet, at the same time, it will allow direct comparison among network properties of individual samples, even in the absence of common taxa among them, as all samples are mapped back to the metacommunity which serves as a normalization step. Thus, these emergent properties can be utilized as universal biomarkers of ecological disturbance.

To exemplify the utility of these metrics of ecological disturbance, we used them to infer the footprint of farming practices at different levels of intensification (conventional, organic and biodynamic) on the fungal community structure in vineyard soils. Our study showed that taxonomic patterns of fungal communities are influenced by geography and climate factors. However, biotic factors, as captured by the local network properties, were crucial at understanding community assembly patterns and the variations derived from the anthropogenic practices, showing global patterns independently of the country ^20^. In this sense, we were able to decipher the different ecological strategies that fungal communities adopt in face of different levels of farming intensification, and this can be used for further explorations on the health status of soils and their response to external stresses (i.e. global change, pathogens invasion, etc). We observed a more generalist-collaborative biota in the soils with less anthropogenic activity (defined by high clustered-low modular networks), and a more niche-specialized biota in those soils with more intensified management (defined by low clustered-high modular networks). We also observed influence of farming practices on the richness of fungal plant pathogens in the soil, where conventional vineyards slightly lower values. However, the role and development of these pathogens in generalists-vs. specialists-based communities should be further studied for estimating the real risk of harbouring plant diseases.

Given the key role that microorganisms play in agri-food systems in general, and in the wine industry in particular, these findings are useful for establishing monitoring programs of crop-associated microbial diversity, supporting the work of alliances such as the global initiative of crop microbiome and sustainable agriculture (https://www.globalsustainableagriculture.org) promoting soil healthiness through agriculture sustainable strategies. We anticipate that this methodological framework could be widely applied to infer ecological disturbances in other natural or anthropic systems, in other fields of interest such as food production or human health, when understanding the effect of practices such as antibiotic and antifungal use, or the microbial origin of healthy vs. pathogenic conditions.

### Box 1.

**Constructing local community networks from metacommunities**

Metacommunity co-occurrence networks have been successfully applied to understand the organization and environmental preferences of microbes ^19^, a fraction of positive trophic interactions ^21^, or defining generalist vs. specialist strategies of particular microbes ^22^. This approach is a common strategy in global or regional studies to give general insights of particular metacommunity structures. However, metacommunity networks do not inform about the actual arrangements of species in local communities, where species may be loosely or densely connected, or display local adaptations to niches or functional guilds. Inferring the local network properties of individual samples characterizes the microbiome of a given sample in terms of its association structure, providing a unique layer of information when studying the biodiversity and stability of a sample, or monitoring its evolution in time and during environmental disturbances. Here, we combine the metacommunity set of associations with local species arrangements to retrieve properties related to particular association arrangements. Considering *n* local communities within a metacommunity, we infer significantly associated pairs of species (both positively and negatively associated). These pairs are later sorted for each local arrangement, so only pairs present in each individual sample are considered, to construct local networks, with particular network properties requiring an ecological interpretation. The ecological interpretation varies according to the positive or negative nature of the associations used to construct the networks, and inferring the emergent properties from these data, may lead to quantifying and understanding the effect of ecological disturbance in natural or anthropic ecosystems. Now we define the most important network properties considered in this study:

#### Connected components

subnetwork in which any two nodes connect to each other by edges, that lack connection to other nodes in the full network.

#### Clustering coefficient

the ratio of triangles and connected triples in the network^23,24^.

#### Average path length

mean of the minimal number of required edges to connect any two nodes^23,24^. **Modularity**: the quality of a partition into **modules** (groups of nodes) using a quantity. A good network partition harbors a higher proportion of edges inside modules compared to the proportion of edges between them^25^.

#### Assortativity

measures the homophyly of the graph, according to node properties or labels (i.e., node degree, which quantifies the number of edges associated to a node). If the coefficient is high, connected nodes tend to have similar values of a given property ^26^.

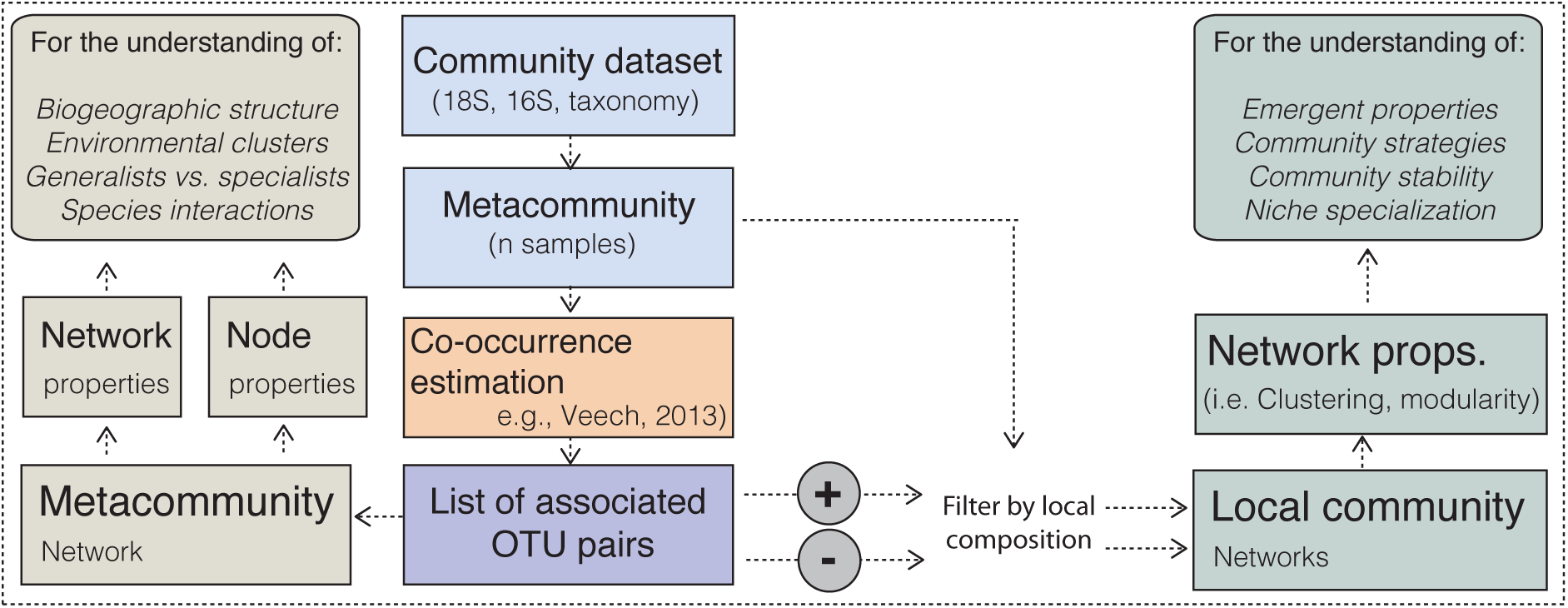

## Results and Discussion

### Fungal community assembly in vineyard soils is affected by biogeography and management factors

Vineyards are human-managed ecosystems and, as one of the most long-lived crops, they are preserved, managed and exploited for centuries in the same soil. Therefore, vineyard soils can be assumed as stabilized and bounded ecosystems, with its biodiversity moulded for decades by the influence of geography, climate, plant-microbe interactions, and farming practices.

Vineyard soils from USA and Spain showed similar alpha- and beta-diversity ranges (**Fig. 1a**), and similar proportions of dominant fungal classes (**Fig. S1**). However, the multivariate ordination of OTU composition showed origin-dependent clusters (**Fig. 1b**), also present in the multivariate ordination of network properties (**Fig. S2**). In a global study from natural ecosystems^27^, fungal communities also exhibited strong biogeographic patterns that appear to be driven by dispersal limitation and climate. Our data partially supports this idea, as the strong geographical distance determined significantly different metacommunities between USA and Spain. Here we also show that the use of different farming practices for vineyard management (conventional, organic and biodynamic) has an impact in fungal community composition (**Fig. 1c,d**), as previously observed by Hartman et al^20^.

**Fig. 1.**
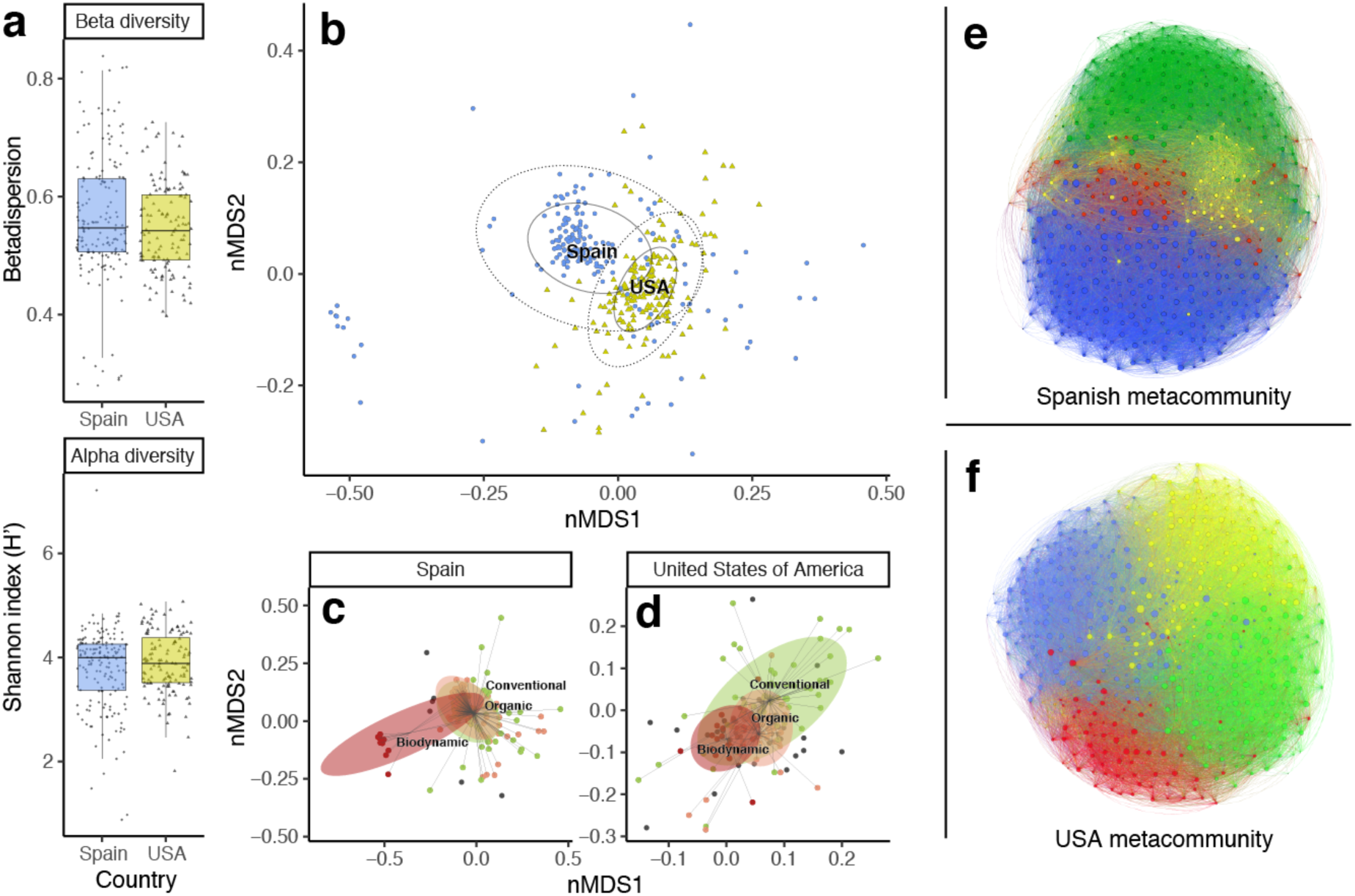
Fungal diversity levels and composition of USA and Spain vineyard soil samples. **a)** comparison of alpha diversity (H’) and beta diversity (betadispersion) of samples between countries; **b)** non-metric multivariate ordination (nMDS) of OTU composition, and definition of country-dependent clusters (anosim r = 0.31, p = 0.001); **c)** multivariate ordination of samples from Spain, and definition of management-dependent clusters (anosim r = 0.18, p = 0.001); **d)** multivariate ordination of samples from USA, and definition of management-dependent clusters (anosim r = 0.17, p = 0.001); **e)** co-occurrence/co-exclusion fungal network of the Spanish metacommunity (fraction with the most abundant OTUs) with coloured modules; **f)** co-occurrence/co-exclusion fungal network of the USA metacommunity (fraction with the most abundant OTUs) with coloured modules.

It is important to highlight that, although Spain has the largest organic grape cultivar worldwide (with 100’000 hectares of organic grapes) ^28^, the fungal diversity of organic vineyards, when compared to conventional vineyards, is actually undetectable **(Fig. 1c)**. However, in the case of USA, with a smaller organic vineyards production, we observed a more evident effect of this type of management on the soil fungal diversity, a half way between conventional and biodynamic managed vineyards **(Fig. 1d)**. Given this different continental effects, both datasets were analysed separately, and two different metacommunities were defined based on co-occurrences/co-exclusions patterns: one for USA samples and one for Spain samples (**Fig. 1e,f**). The study of co-occurrence metacommunity networks, revealed a single connected component with high modularity (ES=0.36, US=0.28), high clustering coefficient (ES=0.31, US=0.57), variable assortativity (ES=0.85, US=0.11), short average path length (ES=1.91, US=1.96), and a higher observed proportion of co-occurrence (ES=0.0696, US=0.0305) and lower co-exclusion (ES=0.0030, US=0.0028) edges out of the total combinations.

### Local network properties explain fungal community composition and structure

Metacommunity networks combined with multivariate techniques and correlative evidence is often used to interpret overall properties of a study system, and although it is a useful approach to catalogue diversity ^29^, to seek for potential associations ^19,30^ and to infer community types within the whole system ^31^, it may be insufficient to advance knowledge on the underlying mechanisms behind some aspects of ecosystem processes or, as in our case, behind different management practices. For the purpose of having a more mechanistic understanding of microbiomes, we evaluated network properties (interpreted as emergent properties) for each local composition of microorganisms individually **(Box 1)**. As expected, local network properties have wider ranges than the overall value of the metacommunity networks of co-occurrences and co-exclusions, for instance modularity (ES=0.003-0.42, US=0.08-0.31), or clustering coefficient (ES=0.44-0.93, US=0.34-0.66) (See full list of ranges in **Table S1**). Despite the geographic differences, most of the network metrics and their interrelationships followed similar trends in the two metacommunities (**Fig. 3**), but also if when merging the two metacommunities in a single global one (see **Fig. 3** footnote). The consistency observed between the USA and Spain metacommunities may indicate that emergent properties of local fungal networks could serve as universal biomarkers of ecological disturbance in soil ecosystems.

**Figure 2.**
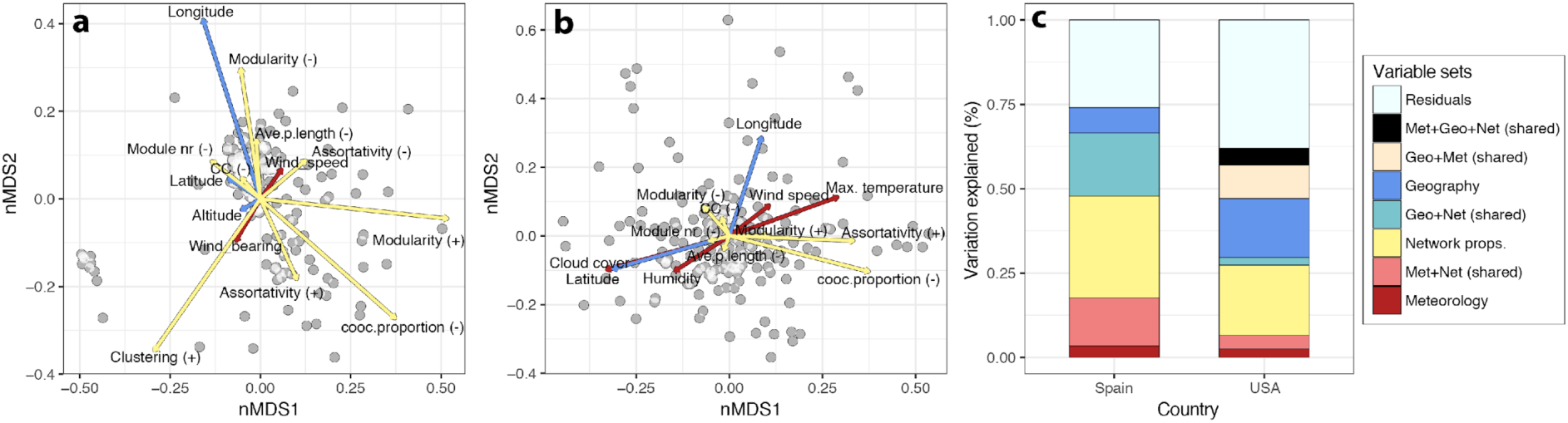
NMDS ordination of soil fungal communities based on Bray-Curtis dissimilarities of OTU composition in (**a**) Spain (stress = 0.17) and (**b**) United States (stress = 0.23). The arrows indicate the direction at which the meteorological factors, geography and network properties fit the best (using envfit function) onto the nMDS ordination space (only shown *p* < 0.01). The size of the arrow is proportional to the strength of the correlation of each variable. The right panel (**c**) shows the percentages of variation in the nMDS ordinations (Spain and USA) explained by the three metadata types through variation partitioning. Not explained (residuals) and shared variation between the three metadata types are also shown.

**Fig. 3.**
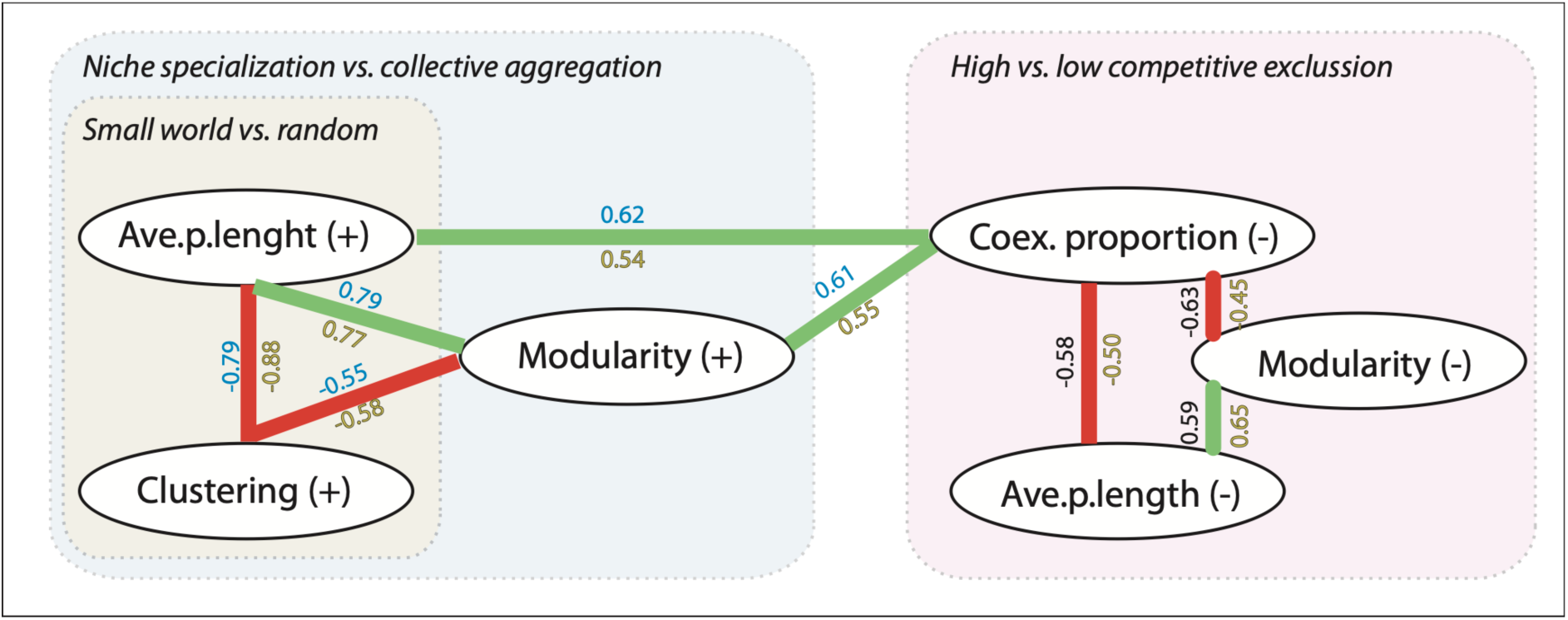
Relationships between network properties. Positive (green) and negative (red) relationships between network properties were obtained based on Spearman’s r correlations > |0.5| in at least one country and *p* < 0.01. Co-occurrences are depicted (+) and co-exclusions (-). Relationships within metacommunities for Spain samples have r values in (blue) and for USA in (yellow). Interpretation of the properties and their associations is indicated in boxes: small-world vs. random networks; niche specialization vs mixed communities; and high vs. low competitive exclusion. Values for relationships between properties in a global metacommunity (merging USA and Spain databases) follow similar trends with the following values (Spearman’s r correlations > |0.5| and *p* < 0.01): Ave.p.lenght(+)--Clustering(+)= −0.88; Ave.p.lenght(+)--Modularity(+)= 0.78; Clustering(+)--Modularity(+)= −0.59; Coex. proportion (-)--Ave.p.lenght(-)= −0.50; Coex. proportion (-)--Modularity(-)= −0.58; Modularity(-)--Ave.p.lenght(-)= 0.74)

To evaluate the relative and combined influence on community assembly of geography, weather and network properties, we conducted a variation partitioning analysis in the two metacommunities. Different community arrangements from USA and Spain were well explained by meteorological factors, geography, and the network properties (from lower to higher variation explanatory power) (**Fig. 2c)**.

In the case of Spain, variability due to weather was explained by an axis of wind speed / altitude; and in the USA an axis of temperature vs. cloud cover (**Fig. 2a**). These were the strongest predictors in an environmental PCA (**Fig. S3**). The effect of geography in each country was also significant and explained a substantial proportion of the community variation. However, in Spain a large proportion of the effect of geography is also shared by network properties (**Fig. 2a**). The remaining unexplained variation (25% in Spain, 33% in USA) could likely be explained by soil organic matter or pH, typical variables reported in a myriad papers^32,33^, which unfortunately we did not measure. However, here we demonstrated that measuring local network properties substantially increased the proportion of community arrangement variations explained by the other abiotic factors (**Fig. 2c)**. These results point out to the crucial relevance of emergent mechanistic processes on community assembly based on species associations, as previously suggested^34^.

### Emergent properties inferred from fungal networks can be used as biomarkers of ecological disturbance in soil samples

To better understand the emergent mechanisms of community structure behind community assembly, we analysed relationships between the network properties and contextualized the ranges of variation into emergent properties from vineyard soils (**Fig. 3**). A Principal Components Analysis (PCA) (**Fig. S2**), showed that the first two axes explained 54% of the variance: the first axis (36%) correlated to positive properties, and the second axis (18%) to negative properties (mostly, the co-exclusion proportion). The inverse relationship in the first axis between clustering coefficient and average path length **(Fig. 3)** indicates the networks can adopt two different structures: a random network (low clustering, high path length) or a small-world structure (high clustering, low path length)^35^. These two properties indicate how densely connected is the network, and thus the degree of activity or interactions between nodes. A densely connected network of co-occurrences may represent organisms preferring the same environmental conditions or be representative of organisms that could have cooperative activities (such as facilitation, syntrophy or/and cross-feeding)^21^. In such small-world networks, random loss of species is unlikely to affect the overall properties of the network, therefore, these are expected to harbour a certain degree of resistance towards perturbations^36^. Furthermore, modularity (+) was inversely associated to the clustering coefficient (+) **(Fig. 3)**. In this context, modularity indicates the degree of separation of the modular components of the networks, and in our results we observe that mycobiomes from vineyard soils can be either densely connected with low niche specialization (generalist), or loosely connected with high niche specialization (specialist) (**Box 2**). A low niche specialization may drive mixed communities, that take advantage of metabolic byproducts by cross-feeding or facilitation ^37^.

On the other hand, competition seems to be central in regulating community assembly over time ^38,39^. Indeed, we observed that highly modular co-occurrence networks, sustain a higher proportion of co-exclusions (**Fig. 3**), which may be quantifying competition processes. It is important to note that co-exclusion related properties, such as modularity (-), average path length (-) or assortativity (-), are estimated by considering pairs of OTUs that occur together less than expected at random. But since these occur, we may interpret them as potential competitors, whose pairs can often coexist ^40^. Indeed, in highly competitive environments, niche partitioning by resource affinity separation or spatial separations, is one of the main strategies that organisms can pursue to survive over time ^38^. In fact, this type of community would harbour metabolic specialist species with reduced overlap^38^, which in the case of fungi may be part of functional guilds that are competing for the same limiting resource through interference competition (such as in mycorrhizal fungi vs. saprotrophs) ^41^. Our results also indicate that more modular specialized communities, with a higher proportion of co-exclusions, harbour lower alpha diversity (H’) (r=-0.41, p<0.001) than densely connected communities. Indeed, competition– colonization tradeoffs can sustain the landscape-scale diversity of microbes that compete for a single limiting resource ^34^. A widely accepted mantra, says that “A higher biodiversity tends to promote a better ecosystem sustainability, through community resistance and resilience promotion”. This statement came not only from the empirical knowledge of soil microbiome by farmers or gut microbiomes by health practitioners, but it is demonstrated by the connection between certain aspects of biodiversity and community structure in the stability of an ecosystem ^42–45^. We argue that in the case of fungi, interference competition and functional specialization are key in regulating soil community diversity^34^, assembly^38^ and the microbiome interaction with vineyards ^46^. Given all these observations, we could state that compositional data combined with a co-occurrence/co-exclusion network approach has allowed us to predict ecological emergent properties such as: i) community stability, ii) niche overlap and iii) fungal competition.

### Farming practices determine fungal ecosystem composition and structure

Our results indicate that management strategies (particularly conventional vs. biodynamic approaches) affect network properties of fungal soil communities, similarly in the two countries studied: USA and Spain. We observed that soils under a biodynamic management had higher clustering coefficient (+), lower modularity (+) and lower co-exclusions proportion than the conventionally managed soils, with organic managed samples tending to show intermediate values between conventional and biodynamic samples (**Fig. 4**) (for full ANOVA results see **Supplementary Table S2)**. This differential action of conventional and biodynamic vineyard management types, with organic practices showing an intermediate effect, has also been recently reported based on soil fertility, nutrient availability, enzyme activity, and earthworm abundance^47^.

**Fig. 4.**
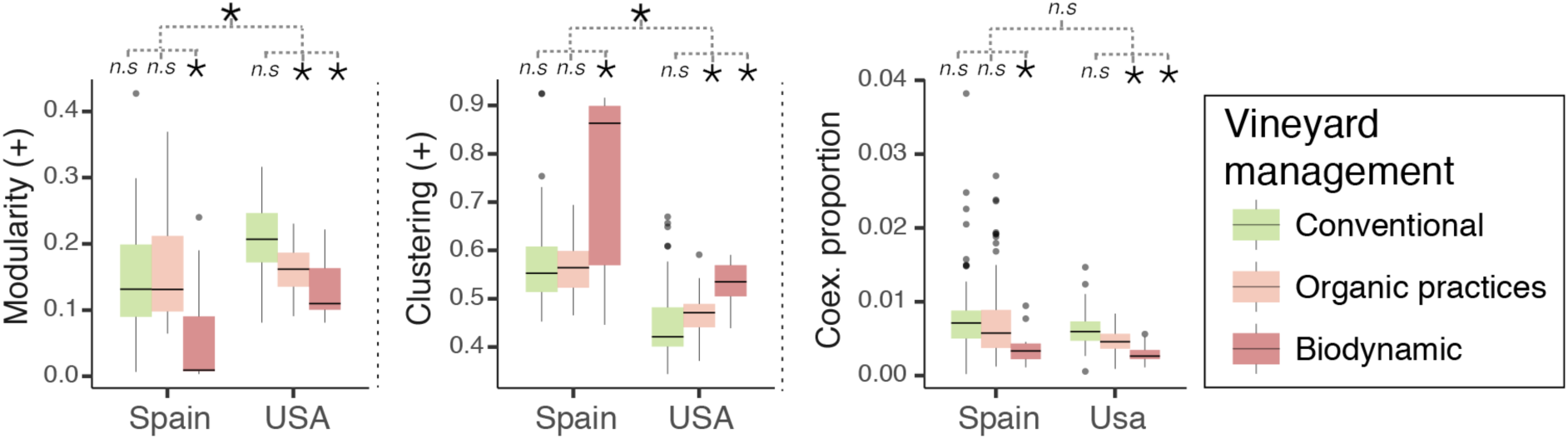
Impact of farming practices in fungal network properties. Boxplot of network properties, (a) Modularity (+), (b) Clustering (+), (c) Co-exclusion proportion (-); under different management practices and countries (Spain-Conventional n=78, Spain-Organic n=79, Spain-Biodynamic n=15; US-Conventional n=65, US-Organic n=39, US-Biodynamic n=20). For each property it is indicated if there is a statistically significant difference, according to a two-way ANOVA (n.s: not significant, *: *p* < 0.01).

In this context, biodynamic-farmed vineyards showed microbial communities closer to: i) small-world networks (higher clustering coefficient (+)); and ii) mixed (generalist-collaborative) communities (lower modularity (+)) (**Fig. 4a,b)**, which are related with enhanced systems homeostasis^38^, reinforcing previous observations of Banerjee et al^48^ in root-associated fungal networks. Conversely, conventionally managed soils gave rise to low clustered, highly modular fungal networks **(Fig. 4a,b)** with a larger proportion of co-exclusions compared to other management types (**Fig. 4c**), reducing, as stated before, the alpha diversity (H’) of the fungal communities. In addition, the use of punctual fertilization programs with high doses of specific nutrients, as in conventional farming, drive a metabolic specialization that may lead to an arrangement of niches^37^, as we observed for conventionally managed samples; in contrast to the more generalist, densely connected communities under biodynamic managements. In parallel, the co-exclusions proportion was associated with lower pathogen richness (r=-0.28, p<0,001) (**Fig. S4a**). Indeed, it is possible to predict that with the lowest co-exclusion proportion values, the probability of having presence of a plant pathogen raises to 80% (**Fig. S4b**). However, it is important to note that a higher richness or abundance of pathogens doesn’t always means a higher risk of disease development. Apart from the environmental conditions, and the direct consequences of phytosanitary programs, the whole community context is also determinant for the vulnerability of the ecosystem, that are likely to be higher in high modular (niche specialized) networks than in those closer to small-world networks^36^. Thus, the community network structure should be taken into account in future in-field studies of ecosystem invasibility by plant pathogens (**Box 2**).

## Conclusions

Ma et al. (2019)^49^ identified the need of systems-level approaches, based on ecological networks, for understanding agro-ecosystems functioning and for studying their sustainability in terms of resilience; for that purpose, the inference of ecological emergent properties^5^ seems a successful strategy to follow. Based on our findings, we can conclude that even in a single ecosystem, human intervention can determine two alternative fungal community assembly strategies: a generalists-based habitat in soils under biodynamic management, or a specialists-based habitat in soils under conventional management. We interpret these situations as alternative equilibrium states of communities (**Box 2**), with those from biodynamic managements closer to small-world networks, potentially related with wild environments. Scaling up to the broader picture, several authors have proposed that two of the major forces driving the current global change in ecosystems functioning - habitat modification and climate change - are expected to select habitat generalists instead of those habitat specialists with lower biodiversity levels and a higher niche partitioning^50,51^. Under this framework **(Box 2)**, our results may lead future theoretical or in-field studies on the biodiversity-stability hypothesis, its relevance for agriculture sustainability, and how human intervention may drive a better future for agro-ecosystems. In addition, the defined ecological emergent properties may be used as biomarkers to measure the effect of farming practices or global change consequences in the health status of soils, but also in other fields such as human health to evaluate the impact of clinical interventions (use of antibiotics/antifungals) or for complement the efforts of connecting human microbiome with health or disease status.

### Box 2.

**Theoretical framework of fungal community structure and functioning in vineyard soils**.

Based in our results, the use of contrasting agricultural management systems (conventional vs. biodynamic) may lead to different emergent properties and community structures in vineyard soils. Specialist vs. generalist habitats appear as the two alternative options that a natural ecosystem could impart, and the level of niche specialization has implications not only on its taxonomic composition, but also on its functionality and in the way that it will respond to external stresses. Following the definition of soil health in agricultural systems given by Kibblewhite et al. (2008)^52^, it can be considered as an “integrative property that reflects the capacity of soil to respond to agricultural intervention, so that it continues to support both the agricultural production and the provision of other ecosystem services”. They also highlight the necessity of optimizing agriculture yields while keeping ecosystem services, as the only way to guarantee the sustainability of the global agriculture system. Thus, the biological sustainability of agro-ecosystems comes from the interaction of the biological processes provided by a diversity of interacting soil organisms and the influence of the abiotic soil environment, with human intervention playing a key role in this interaction.

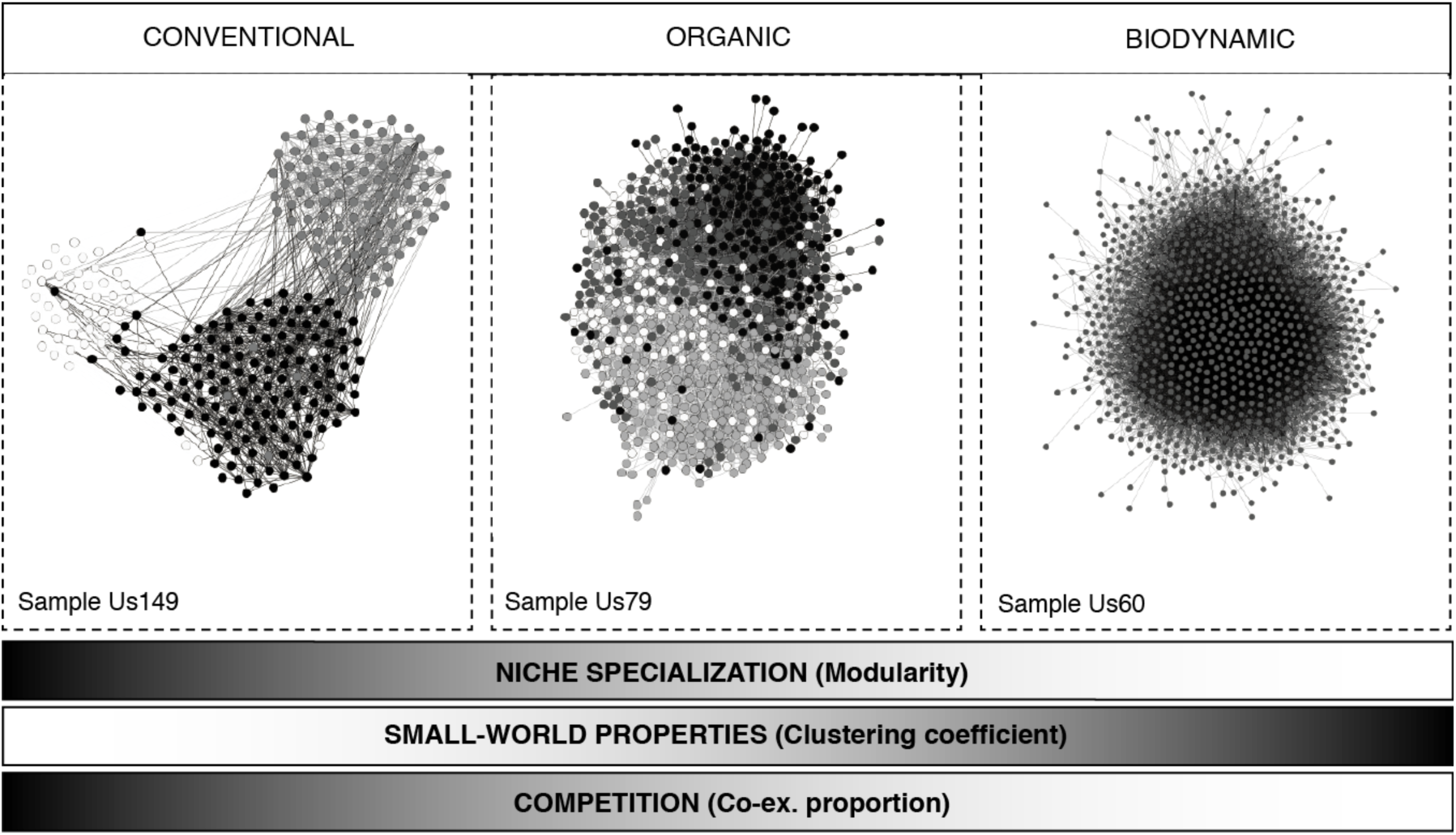

As we report here, the fungal communities favoured under biodynamic management may resemble a community structure close to that in wild cooperation-based environments, as opposed to the highly specialized environment found in conventionally farmed vineyards. As highlighted in a recent consensus paper^53^, the niche specialization found in global soil fungal and bacterial communities and their sensitivity to environmental changes may compromise the future delivery of agro-ecosystem services. This affirmation is based in the demonstrated effect that climate change consequences, such as aridity, have in the reduction in the microbial diversity and abundance of soils. This problem may be higher in highly specialized niche-partitioned environments where functional redundancy and cross-feeding phenomena seems to be lower than in mixed-collaborative systems, where generalist species can be lost with minimal impact on ecosystem processes. Based on that, we can hypothesize that fungal communities that give rise to small-world and collaborative networks, as it is found in biodynamic managed soils, can be more resistant to the continuously changing environment imposed by climate change and land use.

## Methods

### Sample collection, DNA extraction and sequencing

This study is a microbial amplicon-based survey that includes a total of 350 soil samples from vineyards from USA (175 samples; mostly California and southern states) and Spain (175 samples) collected in the period 2015-2018. A general description of the protocol can be delineated as follows: All the samples were of topsoil taken within a depth between 5-10 cm. Each sample from a single block was made pooling together top soil from three random spots in each block and extracting the DNA from this composite sample. Soil samples were stored at −80ºC until DNA extraction. DNA extraction was performed using the DNeasy PowerLyzer PowerSoil Kit (Qiagen). A complete overview of all the samples used in this study and their origin is reported in **Table S4** and in BioProject *PRJNA590645* metadata. Libraries were prepared following the two-step PCR protocol from Illumina and sequenced on an Illumina MiSeq using pair end sequencing (2×300bp). Libraries were prepared by amplifying the 16s rRNA V4 region and the ITS1 region using Biome Makers® custom primers (Patent WO2017096385). Raw files are available under BioProject *PRJNA590645*. Raw sequences were analyzed using Vsearch using default parameters ^54^. Briefly, raw paired-end fastq sequences were merged, filtered by expected error 0.25, dereplicated, and sorted by size. We filtered out chimera sequences and clustered the remaining sequences into 97% identity OTUs, considering in further analyses only groups with at least two sequences. Combined sequences were then mapped to the list of OTUs with at least 97% identity, resulting in an OTU table with OTU sequences quantified per biological sample. OTUs were classified with the SILVA 123 database through the SILVA-NGS pipeline ^55^.

### Sample selection and environmental data

A total of 350 samples were taken from two regions with vineyards: United States of America and Spain. These samples were required to have the following available metadata: geographic location (latitude, longitude and altitude); and climatic information (precipitation intensity, precipitation probability, maximum temperature, minimum temperature, dew point, humidity, environmental pressure, wind speed, wind bearing, wind gust, cloud cover and UV index) obtained from the Dark Sky API site (https://darksky.net/poweredby/). Crop management system (farming practice: conventional, organic or biodynamic) was only available on a subset of samples (124 from USA; 172 from Spain).

### Estimation of emergent properties

For the estimation of network properties, we inferred co-occurrence and co-exclusion probabilities for USA and Spain metacommunities. We checked for a potential effect of not using all the species of the metacommunity by means of Mantel test of Bray Curtis dissimilarities, showing that the filtered communities represent adequately the full local communities. To seek for significant pair-wise patterns of species co-occurrences and co-exclusions, we used a probabilistic method ^56^. This model gives the probability of two species co-occurring or co-excluding each other, at a frequency less or greater than the observed frequency if the two species were distributed independently among sites. The full list of positive and negative significantly associated pairs represents the potential for interactions in the complete metacommunity and/or equivalent environmental distributions. The two lists of positive and negative pairs, were transformed into two species matrices representing the possibility of co-occurrence/co-exclusion in the whole metacommunity. To estimate network properties in each local sample, the two metacommunity-based species matrices were subsequently subset into 350 matrices containing only the species occurring in each of the individual samples. Each of these matrices were transformed into undirected networks using the R package igraph^57^, where nodes represent species, and edges statistically significant co-occurrences/co-exclusions. For each network we estimated the following properties as implemented in igraph: the number of connected components, modularity using the cluster walktrap algorithm ^25^, clustering coefficient defined as average transitivity ^23,57^, average path length^24^ and assortativity^26^ (a global metacommunity network, considering both USA and Spain samples, was punctually used to calculate the relationship between network properties in a unique global context; results reported in the footnote of Figure 3). We also calculated the proportion of co-occurrences and co-exclusions observed, out of the total number of combinations of all the OTUs in the sample. A full representation of the process followed is displayed in **Box 1** (part of this methodology is patent pending (US Patent Application, Serial Number 62947493). All networks were drawn with *gephi* ^58^.

### Statistical analyses

To look for relationships between network properties and plant pathogens, we used Spearman-rank correlations and a principal component analysis (PCA). We assessed if climate or management had an effect on network properties through Spearman correlations and Kruskal-Wallis tests respectively. To estimate the relative contribution of weather, geographic location and network properties in explaining the heterogeneity in the fungal metacommunities, we performed a variation partitioning analysis using the non-metric multidimensional scaling (nMDS) two-dimension scores as the response variables. The three sets of variables were subject to a forward selection procedure, removing collinear variables, prior to their use as explanatory groups of variables ^59^. We further studied if we could predict the presence or absence of plant pathogens (using a curated list of vineyard pathogens), by quantifying the total number of plant pathogens. We further used a model to calculate predicted probabilities of presence of pathogens, by fitting variables (Transitivity (+), Modularity (+), Ave.p.length (-) and co-exclusion proportion) into a generalized linear model (GLM) using a binomial distribution. Statistics were calculated in the R environment using packages *base, vegan*^60^; and drawn in *ggplot2* ^61^.

## Acknowledgements

This work has been partially funded by the Center for Technological and Industrial Development (CDTI) of the Spanish Ministry of Economy, Industry and Competitiveness (MINECO) under the framework of the project IDI-20180120. I.B developed part of this work supported by a MINECO Torres Quevedo Grant (PTQ-16-08253). We thank A. Barberán, C. Cano, D. Almonacid and N. Imam for their comments and discussion, R. Torices and A. Justel for their support in statistical analyses, and A. Ferrero from Biome Makers, Inc. for access to samples and funding.

## Author contributions

R.O., I.B., and A.A. conceived and designed the work; A.A. designed the sampling strategy; R.O., V.J.O., and C.R. contributed to the development of the bioinformatic pipelines; H.O. built the data and computational infrastructure; R.O. performed data analysis; R.O., I.B., and A.A. wrote the paper. All authors reviewed and approved the submitted version.

