## Supplemental material for "Emergent properties in microbiome networks reveal the anthropogenic disturbance of farming practices in vineyard soil fungal communities"

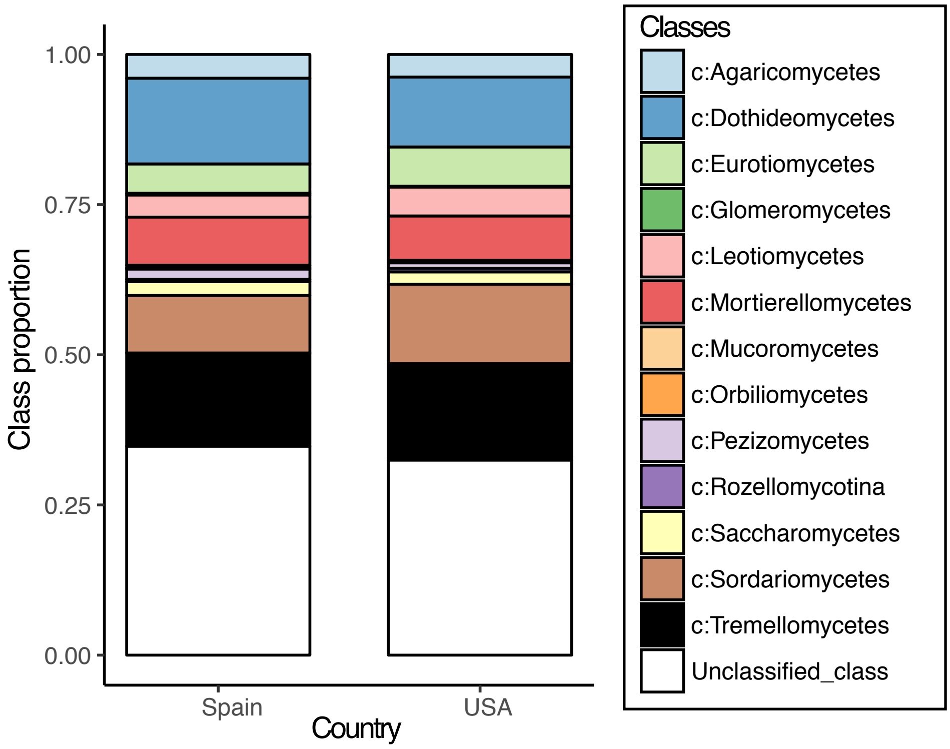


**Figure S1**. Taxonomic profile of major classes of fungi separated by country of origin (Spain and USA), as classified by Unite, and summing the relative abundances of the samples from each country.

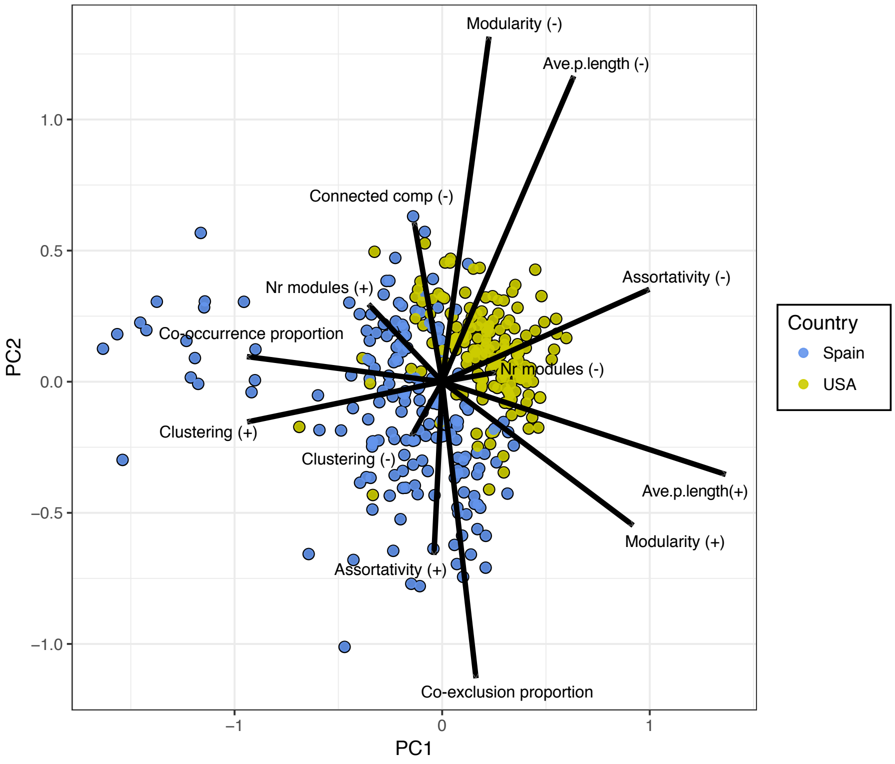


**Figure S2**. PCA of log-transformed scaled network properties separated by Country of procedence (Spain and USA).

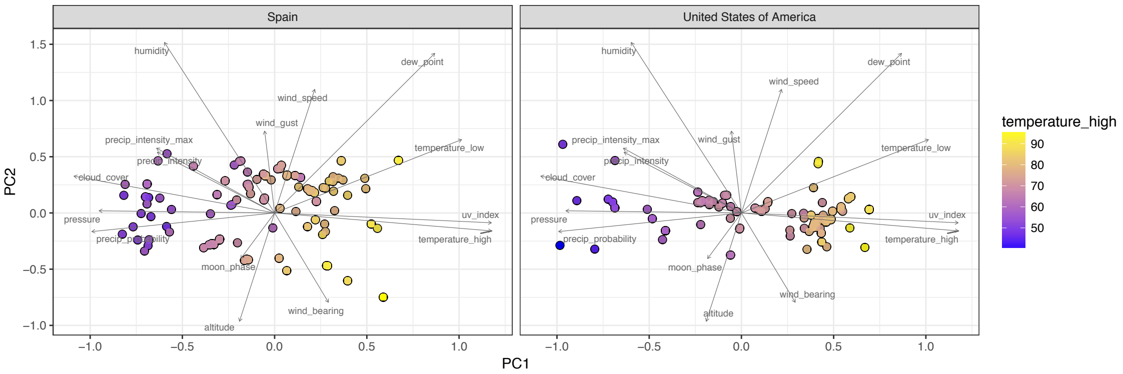


**Figure S3**. PCA of log-transformed scaled environmental variables separated by Country of procedence (Spain and USA).

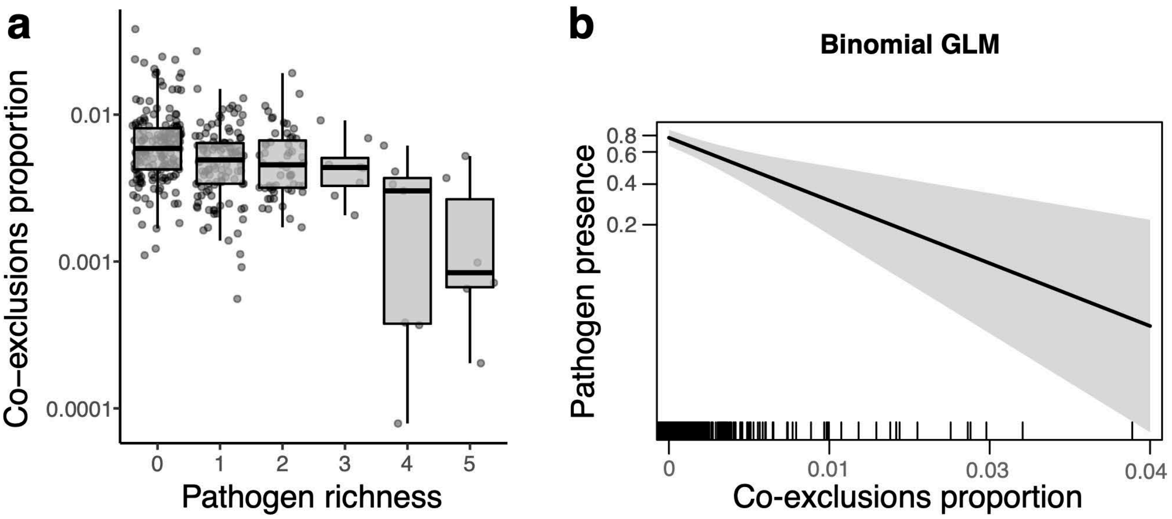


**Figure S4**. Relationship between fungal pathogen richness and co-exclusions proportion of the network. **a)** Boxplot of co-exclusions proportion at different pathogen richnesses (see Table S3 for a detailed list of the vine pathogens considered). **b)** Probability of fungal pathogen presence at different co-exclusions proportion values.

| | **Spain** | | | **USA** | | |
|---|---|---|---|---|---|---|
| **Network property** | **min** | **max** | **mean** | **min** | **max** | **mean** |
| *Co-occurrence proportion* | 0.116 | 0.405 | 0.187 | 0.086 | 0.216 | 0.130 |
| *Clustering (+)* | 0.446 | 0.934 | 0.595 | 0.344 | 0.670 | 0.463 |
| *Modularity (+)* | 0.003 | 0.427 | 0.140 | 0.081 | 0.316 | 0.178 |
| *Nr modules (+)* | 2.000 | 21.000 | 4.846 | 2.000 | 12.000 | 3.817 |
| *Ave.p.length (+)* | 1.189 | 2.134 | 1.643 | 1.574 | 1.889 | 1.755 |
| *Assortativity (+)* | 0.078 | 0.735 | 0.237 | 0.072 | 0.470 | 0.216 |
| *Co-exclusion proportion* | 0.000 | 0.038 | 0.007 | 0.001 | 0.015 | 0.005 |
| *Clustering (-)* | 0.000 | 0.232 | 0.019 | 0.000 | 0.027 | 0.004 |
| *Connected comp (-)* | 1.000 | 6.000 | 1.914 | 1.000 | 9.000 | 1.977 |
| *Modularity (-)* | 0.002 | 0.616 | 0.231 | 0.065 | 0.840 | 0.383 |
| *Nr modules (-)* | 2.000 | 215.000 | 24.417 | 5.000 | 130.000 | 29.543 |
| *Ave.p.length (-)* | 2.000 | 3.891 | 2.935 | 1.231 | 4.116 | 3.379 |
| *Assortativity (-)* | -0.798 | -0.111 | -0.417 | -0.573 | -0.016 | -0.263 |

**Table S1.** Summary of ranges (min, max, mean) for network properties.

| Variable | ANOVA p | Net. Property |
|---|---|---|
| *Country* | 0.0000 | Assortativity (-) |
| | 0.0000 | Ave.p.length (-) |
| | 0.0000 | Ave.p.length (+) |
| | 0.0001 | Co-exclusion proportion |
| | 0.0000 | Co-occurrence proportion |
| | 0.0011 | Nr modules (+) |
| | 0.0000 | Modularity (-) |
| | 0.0000 | Modularity (+) |
| | 0.0000 | Clustering coeff (-) |
| | 0.0000 | Clustering coeff (+) |
| *interaction CxM* | 0.0048 | Assortativity (-) |
| | 0.0076 | Ave.p.length (-) |
| | 0.0000 | Ave.p.length (+) |
| | 0.0000 | Co-occurrence proportion |
| | 0.0049 | Nr modules (-) |
| | 0.0044 | Nr modules (+) |
| | 0.0035 | Modularity (+) |
| | 0.0001 | Clustering coeff (+) |
| *Management type* | 0.0001 | Assortativity (-) |
| | 0.0001 | Assortativity (+) |
| | 0.0000 | Ave.p.length (+) |
| | 0.0039 | Connected components (-) |
| | 0.0000 | Co-exclusion proportion |
| | 0.0000 | Co-occurrence proportion |
| | 0.0000 | Modularity (+) |
| | 0.0165 | Clustering coeff (-) |
| | 0.0000 | Clustering coeff (+) |

**Table S2**. Significant results from two-way ANOVA (Country x Management type) for network properties

| **Disease** | **Pathogen sp.** | **presence** |
|---|---|---|
| Armillaria root rot | *Armillaria mellea* | present |
| Aspergillus rot | *Aspergillus carbonarius* | present |
| Black foot disease | *Ilyonectria robusta* | present |
| Black foot disease | *Campylocarpon fasciculare* | present |
| Black foot disease | *Ilyonectria liriodendri* | present |
| Black foot disease | *Dactylonectria estremocensis* | present |
| Black foot disease | *Campylocarpon pseudofasciculare* | not present |
| Botryosphaeria dieback | *Botryosphaeria dothidea* | present |
| Botryosphaeria dieback | *Lasiodiplodia missouriana* | present |
| Botryosphaeria dieback | *Neofusicoccum parvum* | present |
| Botryosphaeria dieback | *Neofusicoccum australe* | not present |
| Botrytis bunch rot | *Botrytis cinerea* | not present |
| Esca Complex | *Phaeomoniella chlamydospora* | present |
| Esca Complex | *Phaeoacremonium minimum* | present |
| Esca Complex | *Phaeoacremonium hispanicum* | present |
| Esca Complex | *Fomitiporia aethiopica* | not present |
| Esca Complex | *Phaeoacremonium inflatipes* | not present |
| Esca Complex | *Stereum hirsutum* | not present |
| Eutypa dieback | *Eutypella citricola* | present |
| Eutypa dieback | *Cryptovalsa ampelina* | present |
| Eutypa dieback | *Diatrype stigma* | present |
| Eutypa dieback | *Eutypa lata* | not present |
| Petri disease | *Cadophora luteo-olivacea* | present |
| Phomopsis dieback | *Diaporthe ampelina* | present |
| Verticillium wilt | *Verticillium dahliae* | present |

**Table S3.** Disease and pathogen list considered. Presence of that particular pathogen in our dataset is included.

**Table S4.** Sample and metadata list (country and management, more in BioProject *PRJNA590645*)
